# Polar flagellin glycan metaheterogeneity of *Aeromonas hydrophila* strain ATCC 7966^T^

**DOI:** 10.1101/2024.05.20.594992

**Authors:** Kelly M. Fulton, Elena Mendoza-Barberà, Juan M. Tomás, Susan M. Twine, Jeffrey C. Smith, Susana Merino

## Abstract

Motile pathogens often rely upon flagellar motility as an essential virulence factor and in many species the structural flagellin protein is glycosylated. This post-translational modification has been shown to be necessary for proper folding of the flagellin structural proteins and proper function of the flagellar filament in a number of bacterial species. *Aeromonas hydrophila* is a ubiquitous aquatic pathogen with a constitutively expressed polar flagellum. Using a suite of mass spectrometry techniques, the flagellin FlaA and FlaB structural proteins of *A. hydrophila* strain ATCC 7966^T^ were shown to be glycosylated with significant metaheterogeneity: heterologous glycans were observed with variable site occupancy. The penta- and hexa-saccharide glycan chains contained a previously unreported pseudaminic acid derivative with a mass of 422 Da as the linking sugar, followed in sequence by two hexoses, an *N*-acetylglucosamine derivative, a deoxy *N*-acetylglucosamine derivative, and sometimes an additional *N*-acetylglucosamine.

## 3. INTRODUCTION

*Aeromonas* is a genus of gram-negative rod-shaped bacteria that are facultatively anaerobic and found ubiquitously in the environment. This genus belongs to the *Aeromononadaceae* family, *Aeromonadales* order, and *Gammaproteobacteria* class [1]. The taxonomy of this genus is complex and comprises 36 species, broadly categorized as psychrophilic or mesophilic. Psychrophilic species are non-motile, grow at 22-25 °C, and typically only infect fish. Mesophilic species are motile, grow at 35-37 °C, and cause disease in fish (such as tilapia and catfish) and other aquatic animals (such as amphibians and reptiles), as well as humans [2,3,4].

Mesophilic Aeromonads can be isolated from soil and food products [5,6,7]. However, these waterborne pathogens are most commonly associated with marine and freshwater systems, as well as wastewater and sewage effluent [8,9]. Motile aeromonad septicemia is a common disease in fish, with symptoms ranging from skin ulceration to uncoordinated swimming, and often death. Outbreaks in the commercial aquaculture industry have caused significant mortality and associated economic impact. A series of *Aeromonas hydrophila* outbreaks in Alabama, USA, in 2009, for example, resulted in a loss of more than three million pounds of commercial catfish [10]. Subsequent spread of these outbreaks to neighbouring states is estimated to have cost in excess of $12 million [11]. Since many rely upon fish farming for their livelihoods, preventative antibiotic use has become common practice. However, indiscriminate use of antibiotics in aquatic systems is contributing to increasing antimicrobial resistance (AMR) [12,13] with many isolates now characterized as multidrug resistant (MDR) and extensively drug resistant (XDR) [14,15].

Though primarily pathogens of aquatic species, mesophilic Aeromonads, such as *A. hydrophila,* are zoonotic [1]. Previously considered opportunistic pathogens, only posing a threat to immunocompromised individuals, these bacteria now cause disease in immunocompetent populations as well [7]. Transmission is common via ingestion of contaminated food and water [16] leading to infection of the gastrointestinal tract [17,18]. Contact with contaminated water, mud, or sewage effluent can lead to soft tissue infection, especially through open wounds [19,20,21], with severe cases causing necrotizing fasciitis [16,22]. Infections of the blood, eyes, respiratory tract, and urinary tract are also possible [23]. Increasing reports of *Aeromonas* infections in humans and the rise of XDR strains is a concerning combination pointing to an emerging pathogen of clinical and environmental concern.

Pathogenesis is driven by virulence factors that permit bacterial species to maneuver within, adhere to, colonize, and damage a host organism [24]. Flagellar-mediated motility is a common virulence factor for many human pathogens, and glycosylation of the flagellin structural proteins is often a prerequisite for proper structure and function of the flagellar filament [25]. *A. hydrophila* strain ATCC 7966^T^ expresses a polar flagellum, but glycosylation of the structural flagellin proteins has not previously been identified or characterized. This study revealed that polar flagellin proteins FlaA and FlaB were glycosylated at three and five sites, respectively, with highly heterologous glycans. The most frequently observed glycans were pentasaccharides, though a hexasaccharide was additionally observed at a subset of modification sites. Despite the heterogeneity observed, the glycans had a novel pseudaminic acid (Pse) derivative as the linking sugar, followed in sequence by two hexoses (Hex) residues, a variable *N*-acetylglucosamine (GlcNAc) derivative, a variable deoxy *N*-acetylglucosamine (dGlcNAC), and in the case of the hexasaccharides, an additional GlcNAc.

## 4. MATERIALS AND METHODS

### 4.1 Bacterial strains, plasmids, and growth conditions

Bacterial strains and plasmids used in this study are listed in **Table 1**. *A. hydrophila* strain ATCC 7966^T^ and mutants were grown in tryptic soy broth (TSB) or agar (TSA) at 30 °C. *Escherichia coli* strains were grown on Luria-Bertani (LB) Miller broth and LB Miller agar at 37 °C. When required, chloramphenicol (25 µg/mL), rifampicin (100 µg/mL) and spectinomycin (50 µg/mL) were added to the media.

**Table 1.** Bacterial strains and plasmid used in this study.

| Strain or plasmid | Genotype and/or phenotype | Reference |
| --- | --- | --- |
| **Strains** |  |  |
| ***A. hydrophila*** |  |  |
| ATCC7966<sup>T</sup> | *A. hydrophila* strain ATCC7966<sup>T</sup> wild type | [26] |
| ATCC7966-Rif | ATCC7966<sup>T</sup>, spontaneous Rif<sup>r</sup> | [27] |
| ATCCΔAHA4151/77 | ATCC7966-Rif; ΔAHA_4151/4177 | This work |
| ATCCΔAHA4152/71 | ATCC7966-Rif; ΔAHA_4152/4171 | This work |
| ***E. coli*** |  |  |
| MC1061λpir | *thi thr1 leu6 proA2 his4 argE2 lacY1 galK2 ara14 xyl5 supE44 λ pir* | [28] |
| **Plasmids** |  |  |
| pRK2073 | Helper plasmid, Sp<sup>r</sup> | [28] |
| pDM4 | Suicide plasmid, *pir* dependent with *sacAB* genes, oriR6K, Cm<sup>r</sup>. | [29] |
| pDM-AHA4151/77 | pDM4ΔAHA_4151/4177 of ATCC7966<sup>T</sup>, Cm<sup>r</sup>. | This work |
| pDM-AHA4152/71 | pDM4ΔAHA_4152/4171 of ATCC7966<sup>T</sup>, Cm<sup>r</sup>. | This work |

Rif<sup>r</sup>, rifampicin resistant; Cm<sup>r</sup>, chloramphenicol resistant; Sp<sup>r</sup>, spectinomycin resistant.

### 4.2 DNA techniques and sequence analysis

DNA manipulations were carried out using standard procedures. Restriction endonucleases, T4 DNA ligase and alkaline phosphatase purchased from Invitrogen (Thermo Fisher Scientific, Madrid, Spain). PCR analysis for mutant construction was performed using AccuPrime™ Taq DNA polymerase (Invitrogen). Standard PCR analysis was performed using Dream Tap DNA polymerase (Thermo Fisher Scientific). Plasmid DNA was isolated by GeneJet Plasmid Miniprep Kit (Thermo Fisher Scientific) as recommended by the supplier. Nucleotide sequences were determined using the BigDye Terminator v3.1 cycle Sequencing kit (Applied Biosystem).

Genome sequences were retrieved from the NCBI database and comparative genomics of chromosomal regions was performed using Mauve software (version 20150226) [30]. Deduced amino acid sequences were inspected in the Gene Bank database using the NCBI BLASTP network service. Protein family profiling was performed using the Pfam database at the Sanger Center [31].

### 4.3 Construction of in-frame deletion mutants

The ATCCΔAHA4151/77 and ATCCΔAHA4152/71 mutants were generated by in-frame deletion of *AHA_4151/AHA_4177* and *AHA_4152/AHA_4171* chromosomal regions, respectively, by allelic exchange as described previously [29] using primers listed in **Table 2**. Briefly, deletion of chromosomal regions *AHA_4151* to *AHA_4177* or *AHA_4152* to *AHA_4171* were performed by amplification of DNA regions upstream (fragment AB) of *AHA_4151* or *AHA_4152* and downstream (fragment CD) of *AHA_4177* or *AHA_4171*, respectively, in two sets of asymmetric PCRs. Primer pairs A-4151 and B-4151, and C-4177 and D-4177 amplified a DNA fragment of 771bp upstream of *AHA_4151* and 870bp upstream of *AHA_4177*, respectively. Primer pairs A-4171 and B-4171, and C-4152 and D-4152 amplified a DNA fragment of 559bp upstream of AHA_4171 and 772bp downstream of AHA_4152, respectively. DNA fragment AB-4151 and CD-4177, and AB-4171 and CD-4152 were annealed at their overlapping regions and amplified as a single fragment using primers A-4151 and D-4177, or A-4171 and D-4152, respectively. The AD fusion products were purified, *Bgl*II digested, ligated to the pDM4 vector [29] and electroporated to *E. coli* MC1061 (*λpir*). PDM-AHA4151/77 and PDM-AHA4152/71 recombinant plasmids were selected on chloramphenicol plates at 30 °C. Recombinant plasmids were introduced into rifampicin resistant *A. hydrophila* strain ATCC 7966^T^ (ATCC 7966-Rif) by triparental mating using the *E. coli* MC1061 (λ*pir*) containing the plasmid and the *E. coli* mobilizing strain HB101 with pRK2073. Transconjugants were selected on rifampicin and chloramphenicol plates. After sucrose treatment, rifampicin-resistant and chloramphenicol-sensitive transformants were chosen and deletion of DNA fragments was confirmed by PCR.

**Table 2.**
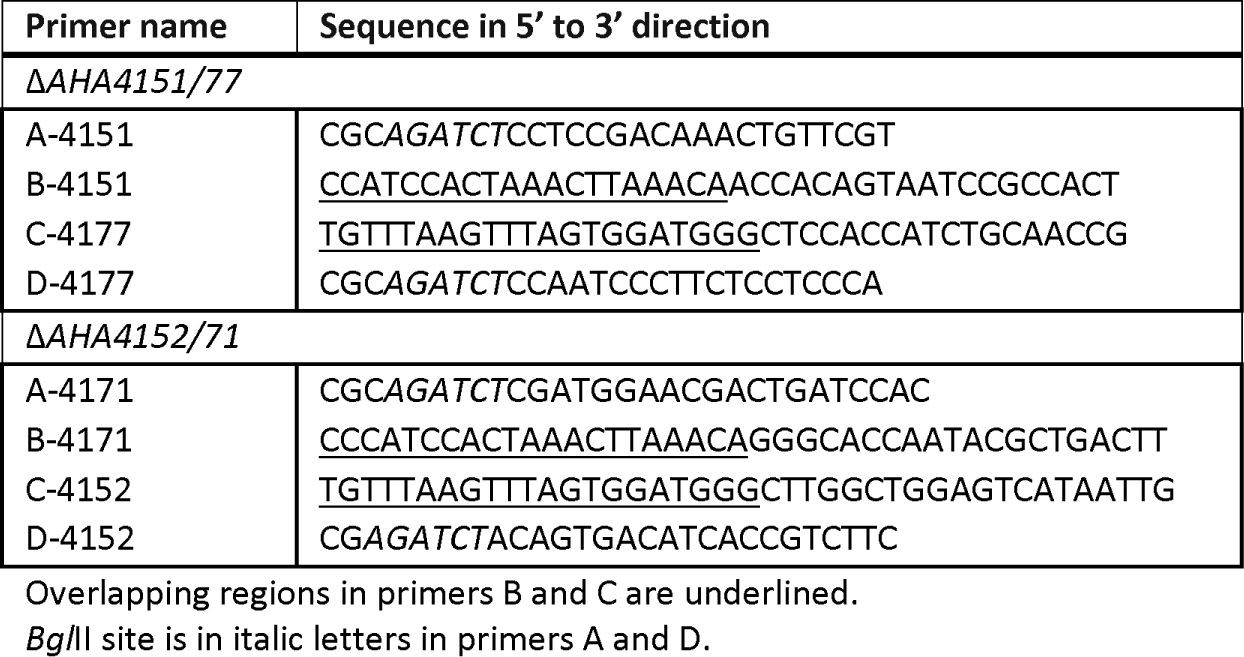
Primers used for the construction of the in frame defined mutants.

### 4.4 Motility assay

Swimming motility was assessed by light microscopy in liquid media and on soft agar plates. Fresh bacterial colonies were transferred with a sterile toothpick into the center of soft agar plates (1% tryptone, 0.5% NaCl, 0.25% agar). Plates were incubated face up for 24 h at 30 °C and motility was assessed by examining bacterial migration through the agar from the center toward the periphery of the plate.

### 4.5 Transmission electron microscopy

Bacterial suspensions were placed on Formvar-coated grids and negative stained with a 2% solution of uranyl acetate (pH 4.1) and observed on a Jeol JEM 1010 (100 Kv with CCD Megaview 1Kx1K) transmission electron microscope (Scientific and Technological service, University of Barcelona, Spain).

### 4.6 Polar Flagellin Purification

Following collection of bacterial cells by centrifugation at 5000 x *g*, polar flagellar filaments were removed from the surface by physical shearing and subsequently purified as described previously [32].

### 4.7 SDS-PAGE and Western Blotting

Purified polar flagellins (2 µg) were separated by 12% SDS-PAGE as described previously [33] and stained with Coomassie Blue (BioRad, Madrid, Spain), according to the manufacturer’s instructions. Western blot analysis of purified flagellins was performed using polyclonal rabbit anti-AH-3 polar flagella antibodies [34], and an alkaline phosphatase-conjugated goat anti-rabbit immunoglobulin G secondary antibody (Sigma-Aldrich, Merck, Barcelona, Spain), both at 1:1000 dilution.

### 4.8 Solution Enzymatic Digest

Purified polar flagellin proteins at a concentration of 0.5 µg/µL in 25 mM ammonium bicarbonate were digested with trypsin (Promega, Madison, WI, USA) using a protein to enzyme ratio of 30:1 (w/w) at 37 °C overnight. Digests were diluted with 0.1% formic acid to 0.15 µg/µL.

### 4.9 Mass Spectrometry Analysis

Peptide digests were analyzed by reversed phase nano liquid chromatography tandem mass spectrometry (nLC-MS/MS). First, 1.5 µg of acidified digest was loaded onto a Phenomonex (Torrance, CA, USA) C18 microtrap (10 x 0.3 mm), followed by analytical separation using a Waters (Milford, MA, USA) BEH130C18 column (100 µm × 10 cm, 1.7 µm particle size, 100 Å pore size) with an UltiMate 3000 LC system (Dionex). Solvent A was 0. 1% (v/v) formic acid in high performance liquid chromatography (HPLC) grade water while solvent B was 0.1% (v/v) formic acid in acetonitrile. With a flow rate of 500 nL/min, the gradient was: 1% to 40% B over 61 min, 40% to 85% B over 3 min, 85% to 1% B over 1 min, and held at 1% B for an 8 min re-equilibration.

MS^1^ and MS^2^ spectra were acquired on a QTOF Ultima (Waters) or an Orbitrap Exploris 480 (Thermo Fisher Scientific). Using the QTOF Ultima, MS spectra were acquired between m/z 200-2000 with MS^2^ fragmentation achieved through collision-induced dissociation (CID). Using the Orbitrap Exploris 480, MS spectra were acquired between m/z 350-1600 with MS^2^ fragmentation achieved through high voltage collision-induced dissociation (HCD) and a normalized collision energy (NCE) of 30%. MS^1^ and MS^2^ resolutions were set to 60,000.

Targeted MS^3^ fragmentation by HCD was performed an Orbitrap Eclipse mass spectrometer (Thermo Fisher Scientific) with a NCE of 20-28% depending on the target ion. Electron transfer dissociation (ETD) was performed on an Orbitrap Eclipse using fluoranthene as the anion donor, a NCE of 35%, and an activation time of 500 ms.

Peaklist (.pkl) files were generated and searched against the NCBI *A. hydrophila* strain ATCC 7966^T^ database using MASCOT search engine (Matrix Science, Boston, MA) for protein identification and sequence coverage [35]. Glycopeptide spectra were interpreted by manual *de novo* sequencing.

## 5. RESULTS

### 5.1 Comparative genomics of the polar flagella glycosylation island (FGI)

Previous bioinformatic analysis of mesophilic *Aeromonas* genomes identified three types of polar FGIs, with *A. hydrophila* strain ATCC 7966^T^ belonging to FGI group II [36]. Comparative genomic analysis of *A. hydrophila* strain ATCC 7966^T^ and *Aeromonas piscicola* strain AH-3 (previously *A. hydrophila* strain AH-3) *fgi* regions showed that only the flanking genes were highly conserved (**Fig 1a)**. The outermost genes, *AHA_4150-4151* and *AHA_4177-4180*, encode proteins orthologous to pseudaminic acid (Pse) biosynthetic proteins, with more than 80% identity to PseBC and PseFGHI of *A. piscicola* strain AH-3, respectively (**Fig 1b**). These *pse* genes were separated by a 27 Kb region containing 11 open reading frames (ORFs) encoding glycosyl-, methyl-, acetyl-, and phospho-transferases, as well as 5 ORFs (*AHA_4172-4176*) encoding proteins involved in fatty acid synthesis, which like Pse biosynthesis proteins, were also highly conserved (**Fig 1b** and **Supplemental Table 1**).

**Fig 1.**
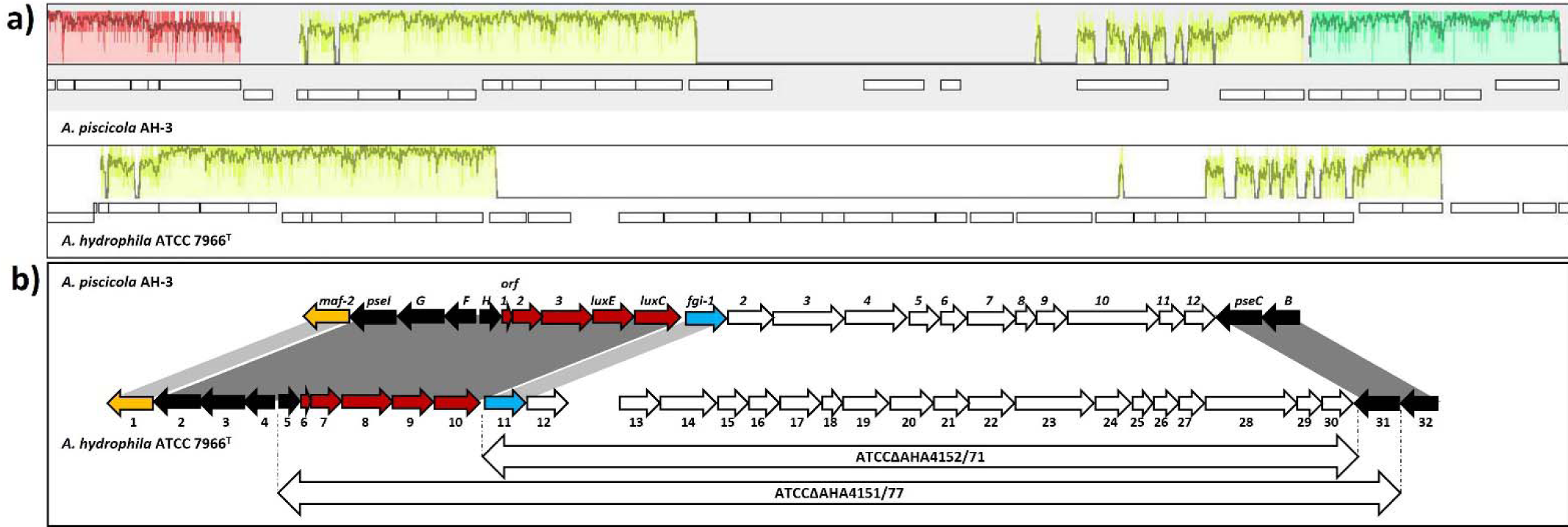
Comparative genomic analysis of *A. hydrophila* strain ATCC 7966^T^ and *A. piscicola* strain AH-3. a) Identification of *A. hydrophila* ATCC 7966^T^ genes involved in polar flagella glycosylation on the chromosomes using progressive Mauve. Yellow fragments in *A. piscicola* strain AH-3 and *A. hydrophila* strain ATCC 7966^T^ indicate homologous segments. b) Schematic comparison of *A. piscicola* strain AH-3 and *A. hydrophila* strain ATCC 7966^T^ polar FGI genes. Sequences with higher than 80% and 55% identity are shown with dark gray and light grey bands, respectively. Black arrows indicate sequences orthologous to *pse* genes. Red arrows indicate sequences orthologous to fatty acid biosynthetic gene. Orange and blue arrows indicate genes orthologous to the motility accessory factor *maf-2* and *fgi-1* of *A. piscicola* AH-3, respectively. White arrows indicate no identity. The long white arrows labelled as ATCCΔAHA4152/71 and ATCCΔAHA4151/77 show the genes removed in these deletion mutants, respectively.

Two in-frame deletion mutants were made to evaluate the role of *fgi* genes in *A. hydrophila* ATCC 7966^T^ polar flagella glycosylation, with deleted genes indicated by the arrows spanning the region between dotted lines in **Fig 1b**. The ATCCΔAHA4152/71 mutant had all genes between *pseC* and *luxC* (including a gene orthogolous to *fgi-1*, a glycosyltransferase in *A. piscicola* strain AH-3) removed. The ATCCΔAHA4151/77 mutant had all genes from and including the *pseC* (*flmB*) to the *pseH* (*flmH*) orthologues removed. ATCCΔAHA4152/71 bacteria showed reduced motility in liquid medium by light microscopy and 40% decreased radial expansion on swimming plates as compared to the wild type strain which showed expected swimming behaviour, while mutant ATCCΔAHA4151/77 was non-motile (**Fig 2a)**. The ATCCΔAHA4151/77 bacteria lacked polar flagellar filaments on the surface, while a fully assembled flagellum was observed on the wild type surface as expected (**Fig 2b**). Western blot analysis of flagellin-enriched fractions revealed that ATCCΔAHA4152/71 bacteria produced flagellin proteins with reduced molecular weight compared to the wild type strain, while mutant ATCCΔAHA4151/77 showed no flagellin reactivity (**Fig 2c**). The observation that glycosylation is important for flagellar assembly and motility prompted in-depth glycoprotein analyses.

**Fig 2.**
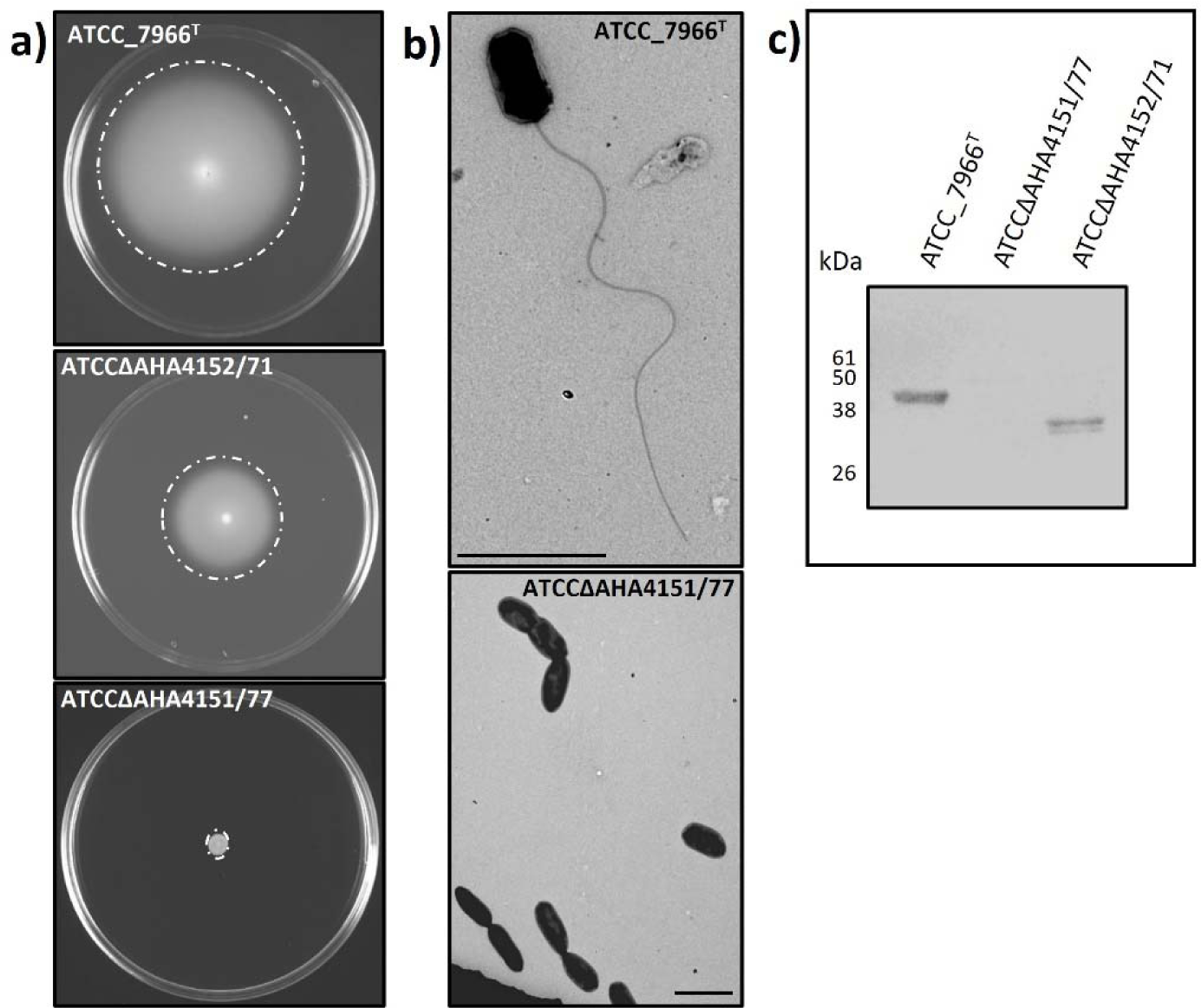
Characterization of FGI region in *A. hydrophila* ATCC 7966^T^. a) Motility of ATCCAHAΔ4152/71 bacteria was reduced compared to wild type bacteria while mutant ATCCAHAΔ4151/77 had no motility at all on soft agar swimming plates. b) Transmission electron microscopy showed a long polar flagellum assembled on the surface of the wild type bacteria with no flagellum present on the surface of mutant ATCCAHAΔ5151/77. Bar = 2 µm. c) Reactivity by Western blot using anti polar flagella antiserum revealed flagellin with reduced molecular weight in the ATCCAHAΔ4152/71 mutant compared to wild type ATCC_7966^T^ and no flagellin present in the ATCCAHAΔ5151/77 mutant.

### 5.2 Glycoprotein Identification

Two genes encode polar flagellin structural proteins within the *A. hydrophila* strain ATCC 7966^T^ genome: *flaA* and *flaB*. Both FlaA and FlaB proteins have a predicted mass of approximately 32 kDa. When separated on SDS-PAGE, both proteins migrated to approximately 45 kDa (**Fig 3a**). The mass difference between the observed migration and the expected protein masses suggested post-translational modification of approximately 12-13 kDa on each protein.

**Fig 3:**
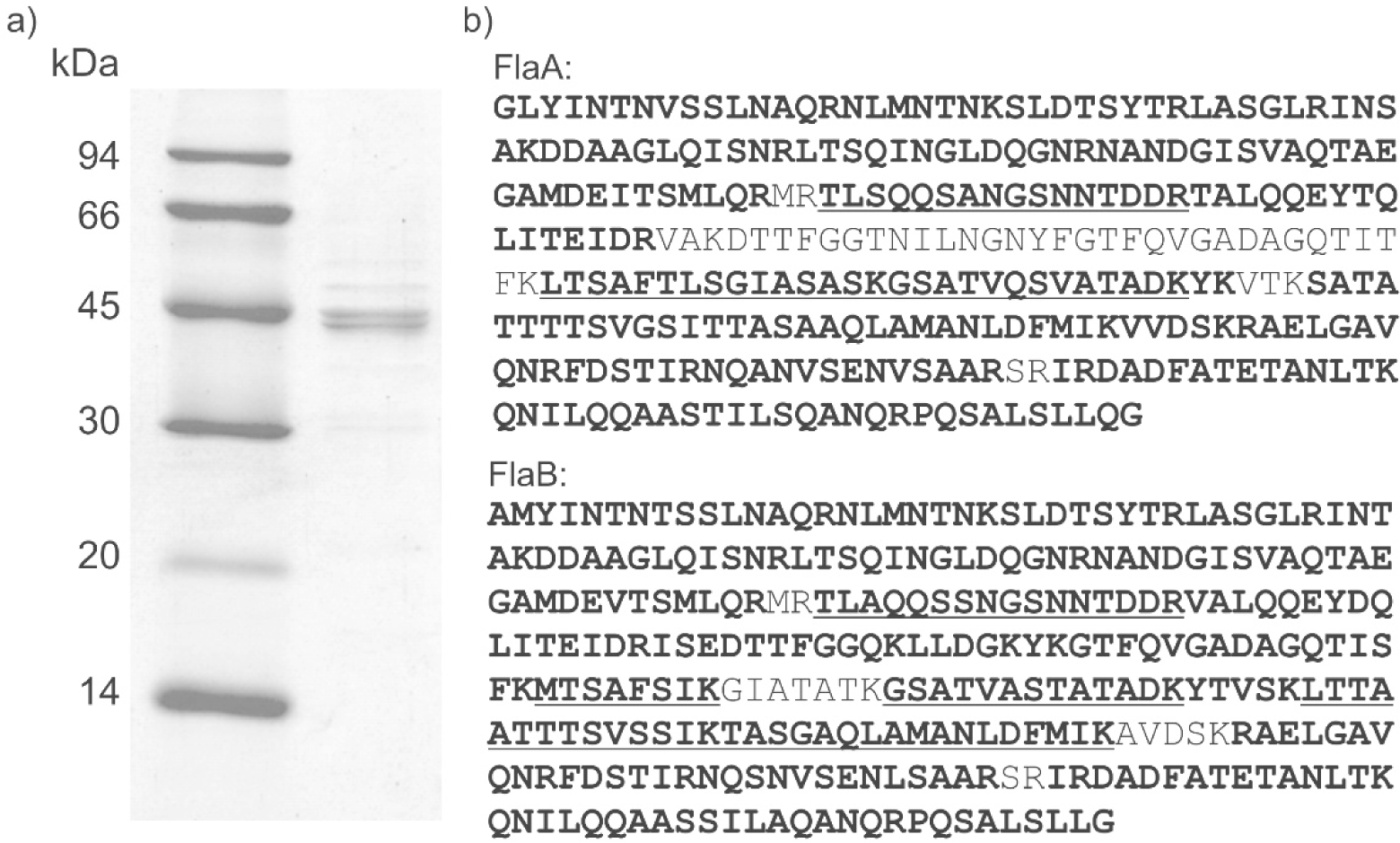
Identifying *Aeromonas hydrophila* ATCC 7966^T^ polar flagellin glycosylation. a) SDS-PAGE of purified polar flagellins revealed that FlaA and FlaB have masses of approximately 45 kDa, despite predicted masses being approximately 32 kDa. b) Sequence coverage of FlaA (86 %) and FlaB (94 %) proteins. Bold text indicates nLC-MS/MS identified tryptic peptides, while underlined sequences are confirmed tryptic glycopeptides.

Polar flagellin protein identities were confirmed by nLC-MS/MS of tryptic digests, with 86% and 94% sequence coverage of FlaA and FlaB, respectively (**Fig 3b**). Several unmatched MS^2^ spectra contained prominent fragment ions in the low m/z region with otherwise poor fragmentation, which is a common observation for glycopeptides.

### 5.3 Glycopeptide Characterization

*De novo* sequencing of unmatched fragmentation spectra revealed that three tryptic FlaA peptides (T^93-108^, T^159-173^, T^174-186^) and five tryptic FlaB peptides (T^93-108^, T^159-166^, T^174-186^, T^192-205^, T^206-222^) were modified with highly heterologous glycans. Glycopeptides FlaA T^93-108^ and FlaB T^93-108^ had the same unmodified mass (1707.5 Da), nearly identical amino acid sequence (TLSQQSANGSNNTDDR and TLAQQSSNGSNNTDDR, respectively), and the same chromatographic retention time. Therefore, the MS^2^ spectra of these isobaric glycopeptides were not resolvable as the m/z values of most of the expected *y* and *b* peptide ions were identical. The MS^2^ spectrum of the unmodified peptide precursor ion at m/z 854.29^2+^ therefore contained *y* and *b* ion series for both peptides (**Fig 4a**). Nonetheless, the glycopeptide spectra associated with these peptide sequences provided the most detailed understanding of the flagellin glycosylation heterogeneity.

**Fig 4:**
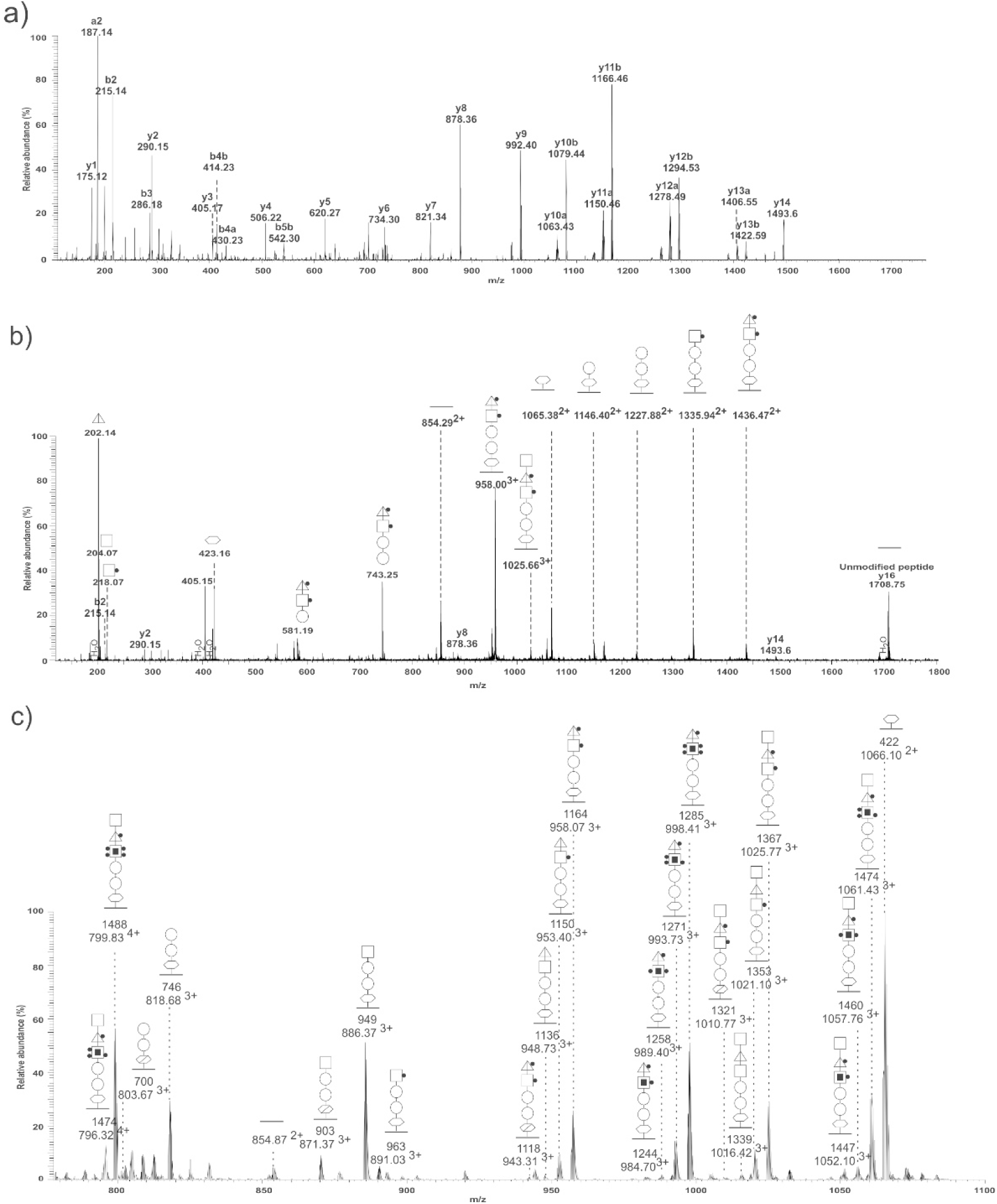
Glycan characterization of an isobaric FlaA and FlaB peptide by nLC-MS/MS. The T^93-108^ peptide of FlaA and FlaB are nearly identical, with an identical unmodified peptide mass of 1707.5 Da, making their spectra unresolvable. Only four *y* and four *b* ions can distinguish these peptides. a) MS/MS spectrum m/z 854.3^2+^ precursor ion produced expected *y* and *b* ion series that confirmed the present of both sequences. b) MS/MS spectrum of the m/z 1025.77^2+^ precursor ion showing the T^93-108^ peptide(s) modified with a representative pentasaccharide glycoform. A doubly charged ion series revealed five neutral losses permitting glycan sequencing of 422-162-162-217-201-203 Da. c) Combined MS^1^ spectrum from 7.5-8.5 min retention time captured the range of glycoform precursor ions associated with the same isobaric peptide sequences. 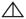: dHexNAc, ◻: HexNAc, ◯: Hex, 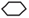: 422 Da pseudaminic acid derivative, 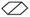: 376 Da pseudaminic acid derivative, ●:Methyl, ◼: Phosphate, 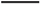: Peptide

Fragmentation of many distinct precursor ions with retention times between 7.5 and 8.5 min resulted in a prominent fragment ion at m/z 1708.75^+^, representing the same unmodified isobaric peptides despite different total masses. The excess mass in the most prominent spectra corresponded to variable glycans with masses ranging from a 422 Da monosaccharide to a 1488 Da hexasaccharide. An unknown glycan oxonium ion at m/z 423.16^+^ and its corresponding dehydrated ion at m/z 405.15^+^ were observed in almost all glycopeptide spectra. Depending on the glycan variant, other oxonium ions at m/z 188.13^+^, 202.14^+^, 204.16^+^, and 218.07^+^ were also observed in the low m/z region in varying combinations. These masses suggested the presence of dHexNAc, Me-dHexNAc, HexNAc, and Me-HexNAc, respectively.

The precursor ion at m/z 1025.77^3+^ had a mass in excess of the 1707.5 Da peptide of 1367 Da. As shown in **Fig 4b**, fragmentation of this glycopeptide produced a clear series of doubly and triply charged ions representing neutral losses from the precursor to the unmodified peptide, permitting glycan sequencing. The mass difference between the ion at m/z 1065.38^2+^ and the unmodified peptide ion at m/z 854.29^2+^ was 422 Da, indicating that this unknown moiety was the linking sugar. Fragment ions at m/z 1146.40^2+^ and 1227.88^2+^ indicated that the second and third sugars were 162 Da Hex residues. In this spectrum, the ion at m/z 1335.94^2+^ revealed a neutral loss of 217 Da, suggesting that the fourth position of the glycan was occupied by Me-HexNAc. However, the monosaccharide mass in the fourth position was highly variable across different spectra associated with the same unmodified peptide ion, ranging from 203 Da to 339 Da. These masses were putatively assigned as HexNAc residues (203 Da) with variable secondary modification of up to one phosphate (+80 Da each) and up to four methyl (+14 Da each) groups. The ion at m/z 1436.47^2+^ indicated that the carbohydrate in the fifth position of the glycan was 201 Da in this spectrum, which was putatively a Me-dHexNAc. However, the neutral loss in the fifth position of the glycan was also variable. In other spectra, a neutral loss of 187 Da suggested that the fifth position of the glycan was variably occupied by either a dHexNAc or a Me-dHexNAc. The mass difference between the precursor and the ion at m/z 1436.47^2+^ was 203 Da, suggesting the terminal sugar in the hexasaccharide glycan was HexNAc. Importantly, only the isobaric FlaA and FlaB peptides were observed to be modified by a hexasaccharide glycan; all other peptides were limited to pentasaccharides.

The combined MS spectrum between 7.5 and 8.5 min was dominated by doubly, triply, and quadruply charged precursor ions representing a range of glycoforms as well as the corresponding unmodified peptide ion for the T^93-108^ FlaA and FlaB sequences at m/z 854.87^2+^ (**Fig 4c**). The precursor ions at m/z 953. 74^3+^, 958.07^3+^, 984.70^3+^, 989.40^3+^, 993.73^3+^, 998.41^3+^, 1021.10^3+^, 1025.77^3+^, 1052.10^3+^, 1057.76^3+^, 1061.43^3+^, and 799.83^4+^ yielded the most prominent pentasaccharide and hexasaccharide glycan masses of 1150.47 Da, 1164.45 Da, 1244.35 Da, 1258.44 Da, 1271.44 Da, 1285.46 Da, 1353 Da, 1367 Da, 1447 Da, 1461 Da, 1474 Da, and 1488 Da, respectively, which were all confirmed by *de novo* sequencing. Peaks corresponding to glycopeptides with intermediate mono-, di-, tri-, and tetra-saccharide glycans were also observed. A few peaks with relatively low ion intensity yielded fragmentation spectra that contained a 376 Da linking sugar (with a glycan oxonium ion at m/z 377.16^1+^) instead of the 422 Da linking sugar observed in all of the prominent glycans.

The remaining six modified peptides had amino acid sequences that were unique to either FlaA or FlaB. None of these peptides were modified by the extended hexasaccharide glycans. They were, however, modified with the same range of mono- to penta-saccharide glycans as were observed on FlaA/Flab T^93-108^. The T^174-186^ peptide sequence, with amino acid sequence GSATVQSVATADK, is shown in **Fig 5** as a representative FlaA glycopeptide harbouring a range of glycans. It had a predicted unmodified peptide mass of 1233.62 Da which was observed at m/z 1234.6^+^ in all spectra along with many expected *y* ions, confirming the peptide identity. These glycopeptides eluted between 10.5 and 11 min. The T^159-166^ peptide sequence, with amino acid sequence MTSAFSIK, is shown in **Fig 6** as a representative FlaB peptide. This peptide had a predicted unmodified mass of 883.45 Da which was observed at m/z 884.50^+^ in all spectra along with many expected *y* ions, confirming the peptide identity. These glycopeptides eluted between 23 and 25.5 min. The mass excess for each glycopeptide shown in both figures corresponded to a prominent pentasaccharide glycan: 1150 Da, 1164 Da, 1243 Da, 1257 Da, 1271 Da, or 1285 Da. All spectra contained the glycan oxonium ions at m/z 423.16^+^ and m/z 405.15^+^, as well as a combination of glycan oxonium ions at m/z 188.13^+^, 202.14^+^, and 218.07^+^, depending on the glycoform. Ions representing partial glycan chains were observed in some spectra at m/z 419.20^+^ (217 Da + 201 Da), m/z 581.31^+^ (162 Da +217 Da +201 Da), m/z 743.31^+^ (162 Da + 162 Da + 217 Da + 201 Da).

**Figure 5:**
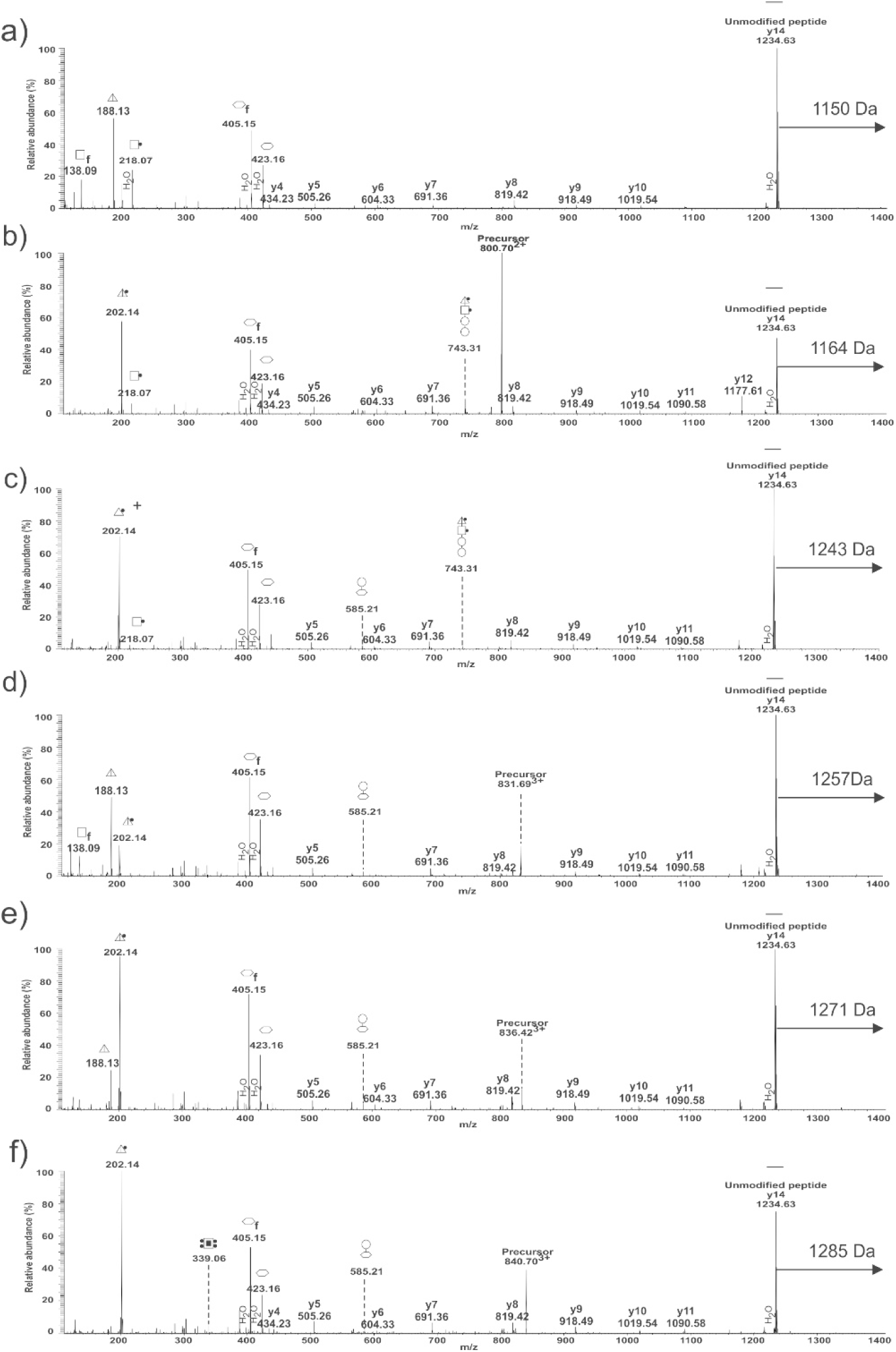
Demonstrating glycan microheterogeneity associated with FlaA with nLC-MS/MS. The FlaA T^174-186^ peptide sequence is GSATVQSVATADK and has a predicted unmodified mass of 1233.62 Da. Fragmentation of precursor ions at m/z 796.03^3+^ (a), 800.71^3+^ (b), 826.29^3+^ (c), 831.69^3+^ (d), 836.02^3+^ (e), and 840.70^3+^ (f) all yielded an ion at m/z 1234.63^+^ and many of the expected *y* ion series for this peptide. The mass difference between the precursors and the unmodified peptide indicated a range of glycan masses (1150.45 Da, 1164.47 Da, 1243.45 Da, 1258.44, 1271.44 and 1285.46 Da, respectively). Glycan oxonium ions and neutral losses are indicated, as well as *y* and *b* ions that confirm the peptide sequence. 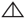: dHexNAc, ◻: HexNAc, ◯: Hex, 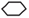: 422 Da pseudaminic acid derivative, ●:Methyl, ◼: Phosphate, 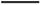:Peptide, f: fragment of associated carbohydrate

**Figure 6:**
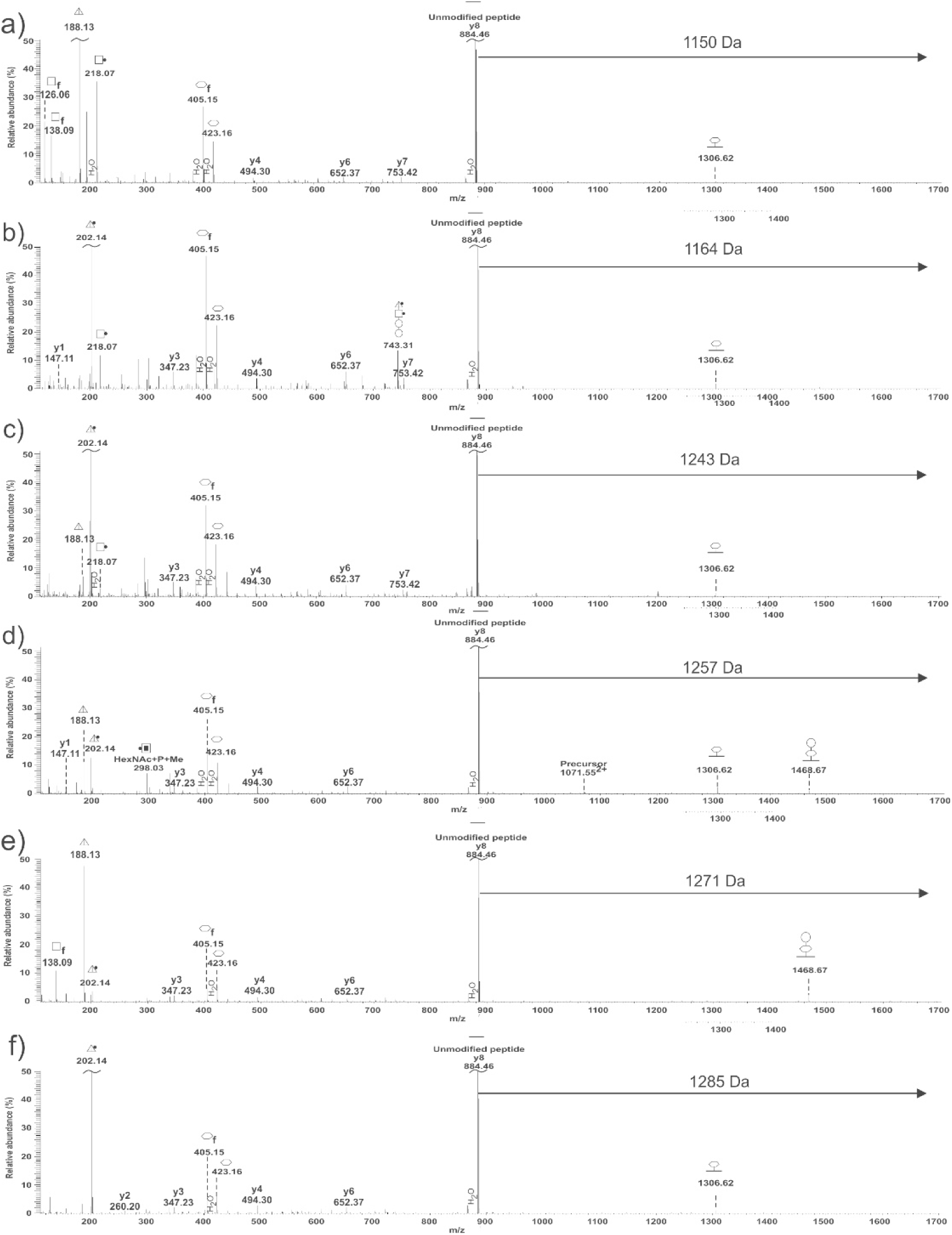
Demonstrating glycan microheterogeneity associated with FlaB with nLC-MS/MS. The FlaB T^160-167^ peptide sequence is MTSAFSIK and had a predicted unmodified mass of 883.45 Da. Fragmentation of precursor ions at m/z 679.30^3+^ (a), 683.98^3+^ (b), 709.97^2+^ (c), 1071.44^2+^ (d), 1078.45^2+^ (e), and 1085.46^2+^ (f) all yielded an ion at m/z 884.46^+^ and many of the expected *y* ion series for this peptide. The mass difference between the precursors and the unmodified peptide indicated a range of glycan masses (1150.45 Da, 1164.47 Da, 1243.45 Da, 1258.44, 1271.44 and 1285.46 Da, respectively). Glycan oxonium ions and neutral losses are indicated, as well as *y* and *b* ions that confirm the peptide sequence. Spectra are zoomed in to 50 % intensity to better visualize low intensity fragment ions. 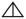: dHexNAc, ◻: HexNAc, ◯: Hex, 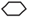: 422 Da pseudaminic acid derivative, ●:Methyl, ◼: Phosphate, 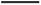: Peptide, f: fragment of associated carbohydrate, 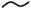: truncated peak

Only three of the eight glycopeptides observed contained an asparagine in the peptide sequence, suggesting that the glycans modifying the polar flagellins were *O*-linked (through serine, threonine or tyrosine) rather than *N*-linked (through asparagine). The precursor ion at m/z 683.98^3+^ corresponded to the FlaB T^160-167^ peptide sequence MTSAFSIK with the 1164 Da glycan modification. It was first fragmented with CID to confirm peptide identity with *y* and *b* peptide fragment ions (**Fig 7a**). Also observed in the CID spectra were glycan oxonium ions at m/z 202.14^+^, 218.07^+^, 405.15^+^, and 423.16^+^ as well as ions representing the entire glycan at m/z 583.00^2+^ and 1164.64^+^. When the same precursor was fragmented by ETD (**Fig 7b**), none of the expected peptide *c* ion series were observed. However, ions corresponding to the expected *c* ion series plus the 1164 Da glycan were observed with respect to *c*_2_ through *c*_8_. This suggested the modification was attached to one of the first two amino acids of the MTSAFSIK peptide. Additionally, *z*_1_, *z*_3_, *z*_4_, *z*_5_, and z_6_ were observed, but *z*_7_ and *z*_8_ were not. Ions corresponding to *z*_7_ and *z*_8_ with the 1164 Da glycan mass were, however, observed at m/z 1901.52^+^ and m/z 1024.52^2+^, respectively. This supported the assignment of Thr161 as the site of glycosylation, which confirmed *O*-linkage.

**Figure 7:**
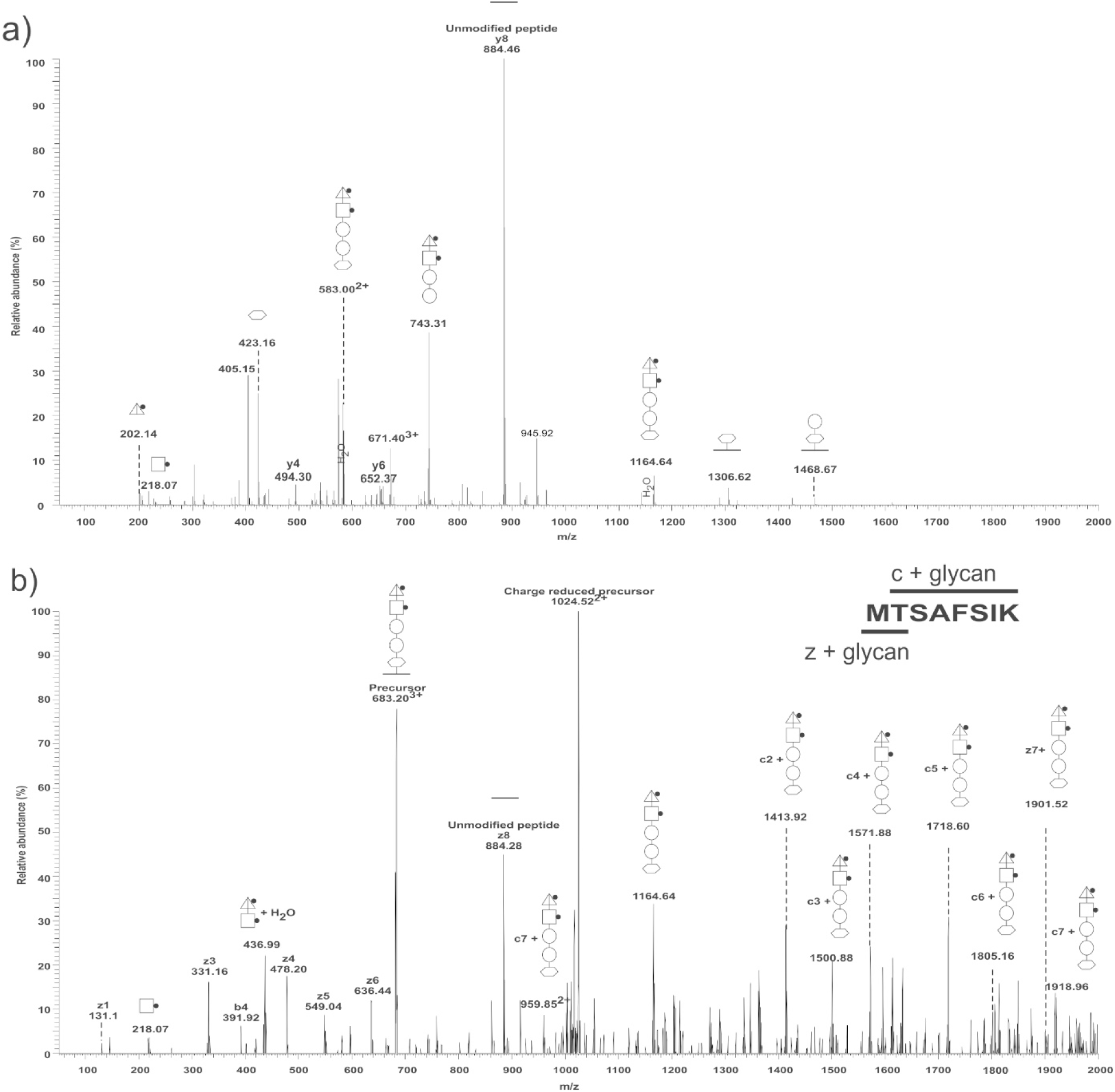
Identification of glycosylation site and linkage by nLC-MS/MS using varied fragmentation methods. a) Collision-induced dissociation (CID) spectrum of the FlaB T^160-167^ MTSAFSIK peptide precursor ion at m/z 683.98^3+^ showing expected *y* and *b* ion series and confirming the amino acid sequence. b) Electron transfer dissociation (ETD) spectrum of the same FlaB T^160-167^ MTSAFSIK peptide precursor ion at m/z 683.98^3+^ produced fragment ions that were 1164 Da greater than *c*2-*c*8 and *z*7-*z*8, indicating that Thr161 was the glycosylation site, and confirmed *O*-linkage. 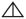: dHexNAc, ◻: HexNAc, ◯: Hex, 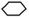: 422 Da pseudaminic acid derivative, ●:Methyl, 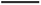: Peptide

### 5.4 Glycan Characterization

Prominent oxonium ions at m/z 204.05^+^ and 188.13^+^ in glycopeptide MS^2^ spectra suggested the presence of HexNAc and dHexNAc, respectively. When the ion at m/z 204.05^+^ was subjected to targeted multistage MS^3^ (**Fig 8a upper panel**), characteristic fragment ions at m/z 84.02^+^, 126.02^+^, 138.09^+^, 144.04^+^, 168.02^+^, and 186.12^+^ were a confirmation that the 203 Da moiety was HexNAc as predicted based on the observed mass. The fragment ion at m/z 138.09^+^ was approximately two-fold more intense than the fragment ion at m/z 144.04^+^, allowing putative assignment of HexNAc as *N*-acetylglucosamine (GlcNAc) and not *N*-acetylgalactosamine (GalNAc) [37]. In a previous study of *A. piscicola* strain AH-3, deletion of *gne* (an epimerase that converts GlcNAc to GalNAc) resulted in reduced polar flagellin molecular weight and motility [38]. However, in *A. hydrophila* strain ATCC 7966^T^, deletion of *gne* had no effect on polar flagellin molecular weight or motility adding support to the putative assignment of GlcNAc to the HexNAC moieties of the glycan.

**Figure 8:**
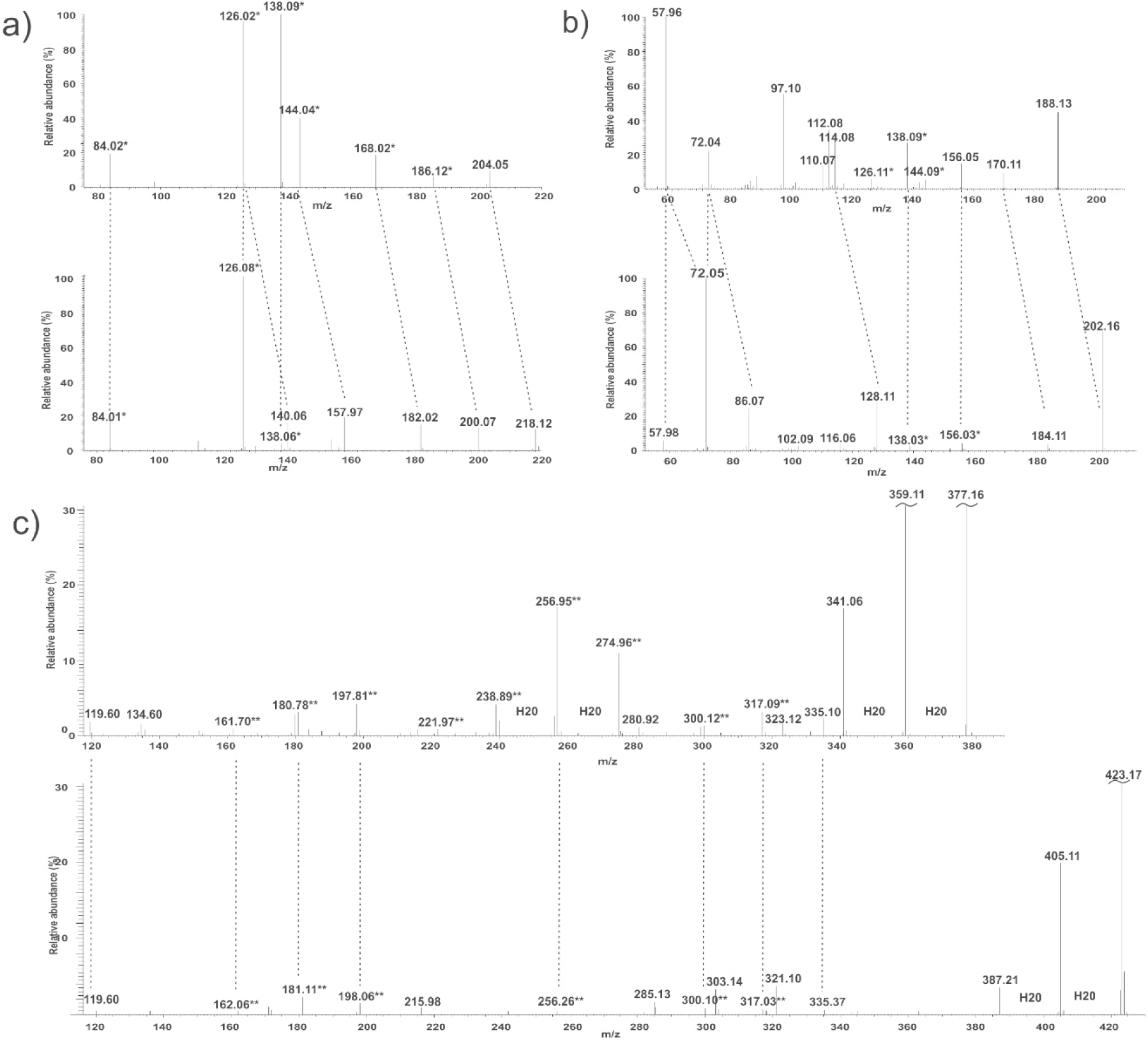
Confirming monosaccharide identities by nLC-MS^3^. a) Targeted MS^3^ of the ion at m/z 204.05^+^ (upper panel) produced fragment ions consistent with an *N*-acetylhexosamine (HexHAc). The intensity of the ion at m/z 138.09^+^ was greater than that of the ion at m/z 144.04^+^, suggesting *N*-acetylglucosamine (GlcNAc) specifically. The MS^3^ fragmentation pattern of the ion at m/z 218.12^+^ (lower panel) revealed a fragmentation pattern similar to that of the ion at m/z 204.05^+^ with a +14 Da mass shift for most peaks, allowing the putative assignment of methylated GlcNAc (MeGlcNAc). b) Targeted MS^3^ of the ion at m/z 188.13^+^ (upper panel) produced fragment ions that were characteristic of HexNAc. In combination with the precursor mass being −16 Da from GlcNAc, this ion was putatively assigned as deoxy *N*-acetylglucosamine (dGlcNAc). Targeted MS^3^ of the ion m/z at 202.16^+^ (lower panel) produced several fragment ions consistent with the dGlcNAc spectrum, as well as several that were shifted by +14 Da, allowing for the assignment of methylated deoxy *N*-acetylglucosamine (Me-dGlcNAc). c) Targeted MS^3^ analysis of the infrequently observed ion at m/z 377.16 (upper panel) and the frequently observed ion m/z 423.17^+^ (lower panel) produced several of the same fragment ions as well as several fragment ions that were consistent with pseudaminic acid (Pse), suggesting both moieties were Pse derivatives. *: characteristic HexNAc fragment ions, **: characteristic Pse fragmentions, 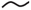: truncated peak.

The variable masses observed in the fourth position of the glycan appeared to correspond to combinations of additional methyl (+14 Da) and phosphate (+80 Da) groups on the GlcNAc. The presence of methyltransferases and phosphotransferases in the *fgi* region (**Supplemental Table 1**) also suggested methylation and phosphorylation were possible. Targeted MS^3^ of the ion at m/z 218.12^+^ (**Figure 8a lower panel**) yielded characteristic GlcNAc fragment ions at m/z 84.01^+^, 126.08^+^ and 138.06^+^ similar to the fragmentation of the ion at m/z 204.05^+^. However, it additionally produced fragment ions at m/z 140.06^+^, 157.97^+^, 182.02^+^, and 200.07^+^, which represented the other observed GlcNAc fragment ions with an additional 14 Da. This allowed for the putative assignment of Me-GlcNAc. MS^3^ of the other variable sugars in position four was not possible, but the mass difference between those neutral losses and GlcNAc, combined with the assignment of Me-GlcNAc to the 217 Da sugar, suggest the other moieties were variably methylated (and phosphorylated) GlcNAc.

Targeted MS^3^ of the ion at m/z 188.13^+^ (**Fig 8b upper panel**) yielded fragment ions at m/z 57.96^+^, 72.04^+^, 97.10^+^, 126.11^+^, 138.09^+^, 144.09^+^, 156.05^+^, and 170.11^+^. Fragments ions at m/z 126.11^+^, 138.09^+^, and 144.09^+^ suggested that despite the reduced mass, this moiety was a derivative of GlcNAc. The mass being 16 Da less than GlcNAc, in combination with the other fragment ions observed having a −16 Da mass shift from other known GlcNAc fragment ions, suggested that the 187 Da moiety was putatively dGlcNAc. Targeted MS^3^ of the ion at m/z 202.16^+^ (**Fig 8b lower panel**) also produced fragment ions at m/z 57.98, 72.04^+^, 138.03^+^, and 156.03^+^, suggesting it was related to the dGlcNAc. However, it additionally had ions at m/z 72.04^+^, 86.07^+^, 128.11^+^, and 184.11^+^ which were 14 Da greater than most of the fragment ions that were different between the two spectra. The 201 Da moiety was therefore putatively assigned as Me-dGlcNAc.

The existence of orthologues to *pse* biosynthesis genes flanking the *fgi* region of *A. hydrophila* strain ATCC 7966^T^, as in *A. piscicola* AH-3, suggested that the glycan would contain a Pse residue. Deletion of *AHA_4151-4177* region that includes *pseC* and *pseH* orthologous genes resulting in loss of flagella formation and motility further associated Pse biosynthesis with the flagellum structure and function. However, there was no glycan oxonium or neutral loss that corresponded directly to Pse (316 Da) in any glycopeptide spectra. In *A. piscicola* strain AH-3, a 376 Da linking sugar was previously shown to be a Pse-like sugar based on similar fragmentation patterns despite the mass difference [32]. In *A. hydrophila* strain ATCC 7966^T^, the linking sugar was most commonly a 422 Da moiety, but a 376 Da moiety was also observed infrequently. Targeted MS^3^ fragmentation of the ion at 377.16^+^ revealed that the 376 Da moiety was the same in both strains, and therefore a Pse derivative (**Fig 8c upper panel**). Furthermore, targeted MS^3^ fragmentation of the ions at 423.15^+^ generated seven fragment ions that were common to both moieties, six of which were known Pse fragment ions (**Fig 8c lower panel**). Though the structural differences accounting for the 106 Da mass difference could not be ascertained, the 422 Da linking sugar was putatively assigned as a novel Pse derivative.

## 6. DISCUSSION

Host-pathogen interactions are dynamic and complex. The severity of infection is dictated by a collection of offensive and defensive mechanisms on both sides of the relationship. Bacteria possess an array of virulence factors, such as flagellar-mediated motility, to enable pathogenesis. Many motile bacteria depend upon glycosylated flagellin proteins for proper flagellum assembly and function. For example, in *Clostridium difficile* strain 630, deletion of the glycosyltransferase gene *0240* resulted in an inability to produce flagellar filaments and reduced motility in a stab agar assay [39]. In hypervirulent strains of *C. difficile*, glycosylation of Type B flagellins was associated with motility as well as adhesion to human Caco-2 intestinal epithelial cells [40]. In *Campylobacter jejuni* strain 81-176 and *Campylobacter coli* strain VC167, deletion of the *Cj1293* glycosyltransferase gene produced bacteria that were non-motile due to a lack of flagellar filament [41]. Furthermore, glycosylation deficient mutants of *C. jejuni* strain 81-176 were defective in autoagglutination, had reduced adherence to human INT407 intestinal cells, and were attenuated in a ferret disease model [42]. Mutation of genes believed to be involved in *Helicobacter pylori* flagellin glycosylation (*HP0178*, *HP0326A*, *HP0326B*, *HP0114*, and *flmD*) resulted in bacteria with reduced or non-existent flagellar filaments and were non-motile [43,44]. Inactivation of gene *rmlB*, which is known to be involved in flagellin glycosylation in *Burkholderia pseudomallei*, resulted in flagella that lacked glycosylation and had reduced motility [45].

The current study confirmed that *A. hydrophila* strain ATCC 7966^T^ glycosylates its polar flagellin proteins, FlaA and FlaB. It does so with a collection of complex oligosaccharides. Despite the observed variability, the linking sugar of glycan was always a Pse derivative attached to the peptide through *O*-linkage. Flagellin glycosylation with a Pse (nonulosonic) or Pse-like linking sugar is common and has been well documented for several bacterial pathogens [46] including *Plesiomonas shigelloides* [47], *C. jejuni* [48], *C. coli* [49], *Clostridium botulinum* [50], and *H. pylori* [43]. Though several structurally related Pse moieties have been observed across those species, the modification was consistently a single monosaccharide. These Pse-like sugars have also been observed as the linking sugar for flagellin glycans in *Aeromonas* species. In *Aeromonas caviae* strain Sch3N, the polar flagellins were modified with a single 316 Da Pse residue at 6-8 sites [51,52] and *A. hydrophila* strain AH-1 flagellins were decorated with a single 403 Da Pse derivative at multiple sites [53]. By contrast, polar flagellin glycosylation in *A. piscicola* strain AH-3 was more complex. In that species, the flagellin glycans were heptasaccharide chains with a 376 Da Pse derivative linking sugar, followed by two Hex, three variable HexNAc derivatives, and an as yet unidentified 102 Da moiety [29]. Regardless of chain length, the conservation of Pse-like sugars in the linking position despite an otherwise varied collection of glycans, suggests it is a critical feature with evidence associating this sugar with proper flagella assembly, motility, adhesion, biofilm formation, and colonization [34,53,54,55,56]. Furthermore, the 422 Da Pse derivative of *A. hydrophila* strain ATCC 7966^T^ observed in this study is also necessary for flagella formation, since deletion of genes encoding PseC and PseH (ATCCΔAHA4151/77 mutant) abolished polar flagella assembly and motility. In contrast, the ATCCΔAHA4152/71 mutant that left the *pse* genes intact produced flagellin with only reduced molecular weight compared to the wild type and reduced but not eliminated motility.

*A. hydrophila* strain ATCC 7966^T^ exhibited flagellin glycan microheterogeneity (variability of glycan composition and configuration). Variable secondary modification of the GlcNAc in position four with up to one phosphate and up to four methyl groups, and variable methylation of the dGlcNAc at position five, combined with variable chain lengths up to a hexasaccharide created the potential for a wide range of glycan permutations, many of which would be isobaric and difficult to resolve. In total, there were seventeen prominent glycan masses observed. Flagellin glycan macroheterogeneity (variability with respect to site occupancy) was also present. Eight distinct sites of modification were observed across the two flagellin proteins through the identification of tryptic glycopeptides, but unmodified versions of the peptides were also present, indicating that site occupancy was not one hundred percent.

Glycan metaheterogeneity (variable micro- and macro-heterogeneity across multiple sites of modification) is a recently coined term in eukaryotes [57] that provides a more accurate description of the observed *A. hydrophila* strain ATCC 7966^T^ flagellin glycosylation patterns. Ten distinct glycan masses, from mono- to penta-saccharide, were observed in association with all known modification sites. There were also seven glycan masses, all hexasaccharides, that were only observed in association with the isobaric FlaA/FlaB peptides. Curiously, these isobaric peptides were in the *N*-terminal D1 domain which contains essential toll-like receptor 5 (TLR5) binding amino acids [58]. This suggested that the variability observed is not random, and nuanced differences likely serve a biological role. By way of precedent, mutation at various glycosylation sites of the *C. jejuni* strain 81-176 FlaA protein produced distinct groups of phenotypes, with three sites leading to reduced motility and truncated flagellar filaments, five sites being defective in autoagglutination, and eleven sites showing no obvious deficiency [59]. However, these findings were not described as glycan metaheterogeneity.

Glycosylation is an essential post-translational modification that enables proper functioning of organisms across all domains of life. It is increasingly recognized in eukaryotes that proper glycosylation involves micro-, macro-, and now meta-heterogeneity. Variation in glycosylation patterns has been shown to affect protein structure and a wide range of biological functions. For example, the absence of fucose specifically at the Asn180 site in the Fc region of human immunoglobulin G increases binding affinity to CD16 and antibody-mediated cellular toxicity [60] while changes to galactose levels at the same site have been implicated in complement-mediated toxicity [61]. Aberrant or uncontrolled glycosylation is also often observed in association with disease states, such as cancers [62], Alzheimer’s disease [63], autoimmunity [64], and a variety of congenital disorders of glycosylation [65].

It is only recently that the terms micro- and macro-heterogeneity have been used to describe glycosylation in prokaryotes, with meta-heterogeneity being used for the first time here to the best of our knowledge. However, there are several bacteria previously shown to have relatively complex glycosylation. For example, 22 distinct glycopeptides were identified during analysis of the *Selenomonas sputigena* flagellin C9LY14 protein [66]. Another example is *C. jejuni* strain NCTC 11168 where up to six different monosaccharide modifications were observed at each site [67]. *Pseudomonas aeruginosa* strain PAK [68] had a glycan that varied from 4-11 sugars. As was already described, *A. piscicola* strain AH-3 had significant glycan variability across multiple sites of modification as well [32]. Even though the application of the term metaheterogeneity is new, it provides an accurate description of the complex *A. hydrophila* strain ATCC 7966^T^ flagellin glycosylation patterns observed. Micro-, macro-, and meta-heterogeneity may provide bacteria an advantage during host-pathogen interactions, possibly helping to safeguard critical virulence factors, such as flagella, against host recognition, host immune responses, and other harsh conditions of the host environment that would threaten to degrade or destroy them. Further evaluation of the FGI is warranted to better understand the complex glycan assembly in *A. hydrophila* strain ATCC 7966^T^.

## 7. ACKNOWLEDGEMENTS

The authors wish to thank Maite Polo for her technical assistance.

## 8. DATA AVAILABILITY STATEMENT

Protein identification and sequence coverage were achieved by searching peaklist files against the publicly available NCBI RefSeq assembly for *Aeromonas hydrophila* subsp. *hydrophila* ATCC 7966^T^ (GCF_000014805.1). Mass spectrometry files generated and analyzed in the current study are available from the corresponding author on reasonable request.

