## SupplementalTable1 for "Polar flagellin glycan metaheterogeneity of *Aeromonas hydrophila* strain ATCC 7966^T^"

**Supplemental Table 1**. *A. hydrophila* strain ATCC 7966^T^ ORFs in the *fgi* region compared to similar genomic region in other *Aeromonas* species

| **ORF** | **Gene** | **Protein size** | **Predicted function** | **Homologous protein** | **% Identity / Similarity** |
| --- | --- | --- | --- | --- | --- |
| 1 | *AHA_4181* | 431 | Motility accessory factor | *A. hydrophila* WP_011707838.1  Maf-2 of *A. piscicola* AH-3 | 100/100  58/70 |
| 2 | *AHA_4180* | 348 | Pseudaminic acid synthase | *A. hydrophila* [WP_259231928.1](https://www.ncbi.nlm.nih.gov/protein/WP_259231928.1?report=genbank&log$=protalign&blast_rank=4&RID=P4SGKUHJ016)  PseI of *A. piscicola* AH-3 | 99/99  94/97 |
| 3 | *AHA_4179* | 418 | UDP-2,4-diacetamido-2,4,6-trideoxy-β-L-altropyranose hydrolase | *A. hydrophila* WP_011707836.1  PseG of *A. piscicola* AH-3 | 100/100  83/87 |
| 4 | *AHA_4178* | 231 | Pseudaminic acid cytidylyltransferase | *A. hydrophila* WP_011707835.1  PseF of *A. piscicola* AH-3 | 100/100  87/94 |
| 5 | *AHA_4177* | 178 | UDP-4-amino-4,6-dideoxy-N-acetyl-β-L-altrosamine N-acetyltransferase | *A. hydrophila* WP_263680050.1  PseH of *A. piscicola* AH-3 | 99/100  92/93 |
| 6 | *AHA_4176* | 77 | Acyl carrier protein | *A. hydrophila* WP_011707833.1  Orf-1 of *A. piscicola* AH-3 | 100/100  95/97 |
| 7 | *AHA_4175* | 249 | 3-oxoacyl-ACP reductase | *A. hydrophila* WP_011707832.1  Orf-2 of *A. piscicola* AH-3 | 100/100  91/93 |
| 8 | *AHA_4174* | 456 | AMP-dependent synthase | *A. hydrophila* WP_011707831.1  Orf-3 of *A. piscicola* AH-3 | 100/100  84/89 |
| 9 | *AHA_4173* | 352 | Acyl protein synthase | *A. hydrophila* WP_011707830.1  LuxE of *A. piscicola* AH-3 | 100/100  90/93 |
| 10 | *AHA_4172* | 395 | Acyl-CoA reductase | *A. hydrophila* WP_011707829.1  LuxC of *A. piscicola* AH-3 | 100/100  83/87 |
| 11 | *AHA_4171* | 318 | Glycosyltransferase family 2 | *A. hydrophila* WP_164927770.1  Fgi-1 of *A. piscicola* AH-3 | 100/100  55/70 |
| 12 | *AHA_4170* | 366 | Glycosyltransferase family 8 | *A. hydrophila* WP_011707827.1  *A. dhakensis* WP_005306709.1 | 100/100  98/99 |
| 13 | *AHA_4169* | 378 | Glycosyltransferase family 1 | *A. hydrophila* WP_237701933.1  *A. dhakensis* WP_231504425.1 | 100/100  98/99 |
| 14 | *AHA_4168* | 434 | Nucleotide sugar dehydrogenase | *A. hydrophila* WP_011707825.1  *A. dhakensis* WP_041202566.1 | 100/100  99/99 |
| 15 | *AHA_4167* | 277 | Glycosyltransferase family 2 | *A. hydrophila* WP_011707824.1  *A. dhakensis* WP_095591648.1 | 100/100  97/98 |
| 16 | *AHA_4166* | 257 | Glucose-1-phosphate cytidylyltransferase | *A. hydrophila* WP_011707823.1  *A. dhakensis* WP_206220935.1 | 100/100  85/93 |
| 17 | *AHA_4165* | 355 | CDP-glucose 4,6-dehydratase | *A. hydrophila* WP_164927769.1  *A. dhakensis* WP_005306694.1 | 100/100  97/97 |
| 18 | *AHA_4164* | 187 | dTDP-4-dehydrorhamnose 3,5-epimerase | *A. hydrophila* WP_011707821.1  *A. dhakensis* WP_005306691.1 | 100/100  97/98 |
| 19 | *AHA_4163* | 407 | Methyltransferase | *A. hydrophila* WP_011707820.1  *A. dhakensis* WP_005306688.1 | 100/100  98/98 |
| 20 | *AHA_4162* | 368 | Aminotransferase | *A. hydrophila* WP_011707819.1  *A. dhakensis* WP_005306686.1 | 100/100  98/99 |
| 21 | *AHA_4161* | 259 | AdoMet-dependent methyltransferase | *A. hydrophila* WP_085734468.1  *A. veronii* WP_245098543.1 | 99/99  94/97 |
| 22 | *AHA_4160* | 359 | SAM-dependent methyltransferase | *A. hydrophila* WP_011707818.1  *A. dhakensis* WP_005306680.1 | 100/100  95/96 |
| 23 | *AHA_4159* | 631 | Hypothetical protein | *A. hydrophila* WP_011707817.1  *A. dhakensis* WP_043172348.1 | 100/100  97/97 |
| 24 | *AHA_4158* | 318 | Hypothetical protein | *A. hydrophila* WP_011707816.1  *A. dhakensis* WP_005306674.1 | 100/100  96/97 |
| 25 | *AHA_4157* | 173 | Adenylyl-sulphate kinase | *A. hydrophila* WP_011707815.1  Fgi-8 of *A. piscicola* AH-3 | 100/100  41/58 |
| 26 | *AHA_4156* | 191 | Hypothetical protein | *A. hydrophila* WP_011707814.1  *A. dhakensis* WP_005306671.1 | 100/100  98/98 |
| 27 | *AHA_4155* | 240 | Formyltransferase | *A. hydrophila* WP_011707813.1  *A. dhakensis* WP_095591644.1 | 100/100  98/99 |
| 28 | *AHA_4154* | 798 | Phosphoenolpyruvate synthase | *A. hydrophila* [WP_011707812.1](https://www.ncbi.nlm.nih.gov/protein/WP_011707812.1?report=genbank&log$=protalign&blast_rank=1&RID=P1EPH88N016)  Fgi-10 of *A. piscicola* AH-3 | 100/100  35/50 |
| 29 | *AHA_4153* | 212 | Glutamine amidotransferase | *A. hydrophila* [WP_041216839.1](https://www.ncbi.nlm.nih.gov/protein/WP_041216839.1?report=genbank&log$=protalign&blast_rank=2&RID=P1ENZ83Y013)  Fgi-11 of *A. piscicola* AH-3 | 100/100  35/45 |
| 30 | *AHA_4152* | 251 | Nucleotydyl transferase | *A. hydrophila* [WP_011707810.1](https://www.ncbi.nlm.nih.gov/protein/WP_011707810.1?report=genbank&log$=protalign&blast_rank=1&RID=P1DV7D88013)  Fgi-12 of *A. piscicola* AH-3 | 100/100  44/56 |
| 31 | *AHA_4151* | 377 | UDP-4-amino-4,6-dideoxy-N-acetyl-*β*-L-altrosamine transaminase | *A. hydrophila* [WP_011707809.1](https://www.ncbi.nlm.nih.gov/protein/WP_011707809.1?report=genbank&log$=protalign&blast_rank=1&RID=P1E6U0ET016)  PseC of *A. piscicola* AH-3 | 100/100  81/88 |
| 32 | *AHA_4150* | 334 | UDP-N-acetylglucosamine 4,6-dehydratase | *A. hydrophila* [WP_011707808.1](https://www.ncbi.nlm.nih.gov/protein/WP_011707808.1?report=genbank&log$=protalign&blast_rank=1&RID=P1EDSSVH016)  PseB of *A. piscicola* AH-3 | 100/100  96/96 |
